# Long-range transport of biomolecular condensates by Marangoni effect

**DOI:** 10.64898/2026.09.25.753395

**Authors:** Qi Guan, Xiaoli Ma, Xin Guan, Jizhong Lou, Hong Zhang, Zhi Qi

## Abstract

How mesoscale cargos such as biomolecular condensates are transported and spatially positioned within cells remains a fundamental question in cell biology. During Caenorhabditis elegans zygote polarization, a substantial fraction of P granules undergoes persistent anterior-to-posterior motion that cannot be fully explained by dissolution–recondensation. Here, combining microfluidic reconstitution, quantitative micropipette aspiration and live-cell imaging, we show that a MEG-3 concentration gradient generates interfacial-tension asymmetry and drives the directed transport of PGL-3–RNA co-condensates in vitro and P granules in vivo through the Marangoni effect. This framework enables parameter-free quantitative predictions of condensate transport. Orthogonal molecular perturbations abolish or reverse directional migration, directly linking interfacial properties to transport behavior and establishing the specificity of this mechanism. Together, our findings identify the Marangoni effect as a previously unrecognized mechanism for intracellular transport of biomolecular condensates and provide direct causal evidence that condensate emergent properties can contribute to physiological organization.

## Introduction

The establishment of cell polarity is a fundamental symmetry-breaking process that requires the precise spatiotemporal organization of intracellular components (*1, 2*). In *Caenorhabditis elegans* embryos, polarity cues orchestrate the partition of P granules towards the posterior. These RNA-rich biomolecular condensates contain RNA-binding protein like PGL-1, PGL-3 and associated RNAs (*3, 4*). The redistribution of P granules is essential for germline specification and RNA-based epigenetic inheritance (*2, 3, 5, 6*).

This phenomenon has been explained by several well-established mechanisms, including bulk cytoplasmic advection (*7–9*) and position-dependent dissolution– recondensation regulated by opposing anterior–posterior gradients of the RNA-binding proteins MEX-5/6 (*3, 10–12*). However, these population-level mechanism does not explain the persistent, directed displacement of individual granules over tens of micrometers during several minutes’ polarization window. This distinction is supported by studies on MEG-3, a protein forms an anterior-to-posterior gradient opposing MEX-5/6 (*13, 14*) and accumulates at the P granule interface to reduce its interfacial tension through interaction with PGL-3 and formation of Pickering-like interfacial stabilization layer (*15, 16*). An interaction-attenuated MEG-3_HMGL−_ mutant, whose four conserved residues in the HMG-like motif were substituted with alanine (F705A, Y708A, Y713A, and N725A), preserves bulk cytoplasmic streaming and dissolution–recondensation, but abolishes P granule polarization (*17*). This result indicates that interfacial regulation contributes to P granule positioning. Moreover, time-lapse imaging shows that a substantial fraction ∼40% of anterior P granules move persistently towards the posterior (*3, 8*) (Supplementary Fig. 1a), traveling ∼50 μm within ∼500 seconds. Such rapid, directional transport is difficult to reconcile with dissolution–recondensation alone and points to a missing mechanism for transporting intact condensates.

Given that MEG-3 lowers the interfacial tension of PGL-3 condensates, we hypothesized that its anterior-to-posterior concentration gradient establishes a corresponding interfacial tension gradient across P granule interfaces. This scenario closely resembles the classical Marangoni effect in interfacial science and fluid mechanics (*18, 19*), in which interfacial tension gradients generate tangential interfacial flows that propel fluid droplets toward regions of lower interfacial tension. Although Marangoni-driven motility has recently been demonstrated in synthetic aqueous phase-separated droplets (*20, 21*), whether endogenous biochemical gradients similarly harness this mechanism to transport functional biomolecular condensates remains unknown. More broadly, condensates pose a transport problem on efficiency and specificity that conventional mechanisms do not fully solve. They are large enough to diffuse slowly, lack membranes or dedicated motor adaptors, and experience strong, size-dependent resistance in the crowded poro-viscoelastic cytoplasm. (*22, 23*). The physical principles that enable rapid, long-range and directional transport therefore remain unclear (*22, 24, 25*).

In this study, we combined microfluidic reconstitution, micropipette aspiration, live-cell imaging, and quantitative particle tracking to demonstrate that MEG-3 reduces the interfacial tension of PGL-3–RNA co-condensates, while a MEG-3 concentration gradient drives their directed migration toward regions of higher MEG-3 concentration. Condensate migration velocity increased with condensate size and quantitatively matched the parameter-free predictions of the Marangoni effect. Weakening the MEG-3–PGL-3 interaction with the MEG-3_HMGL-_ mutant abolished both interfacial-tension regulation and directional transport, whereas the endogenous isoform MEG-3 Δ16 generated by a 48 nucleotides (48-nt) alternative-splicing event that removes 16 amino acids (aa 190-205) from its intrinsically disordered region reversed both responses. In *Caenorhabditis elegans* embryos, the P granule motion showed the same size dependence predicted from independently measured material parameters. Finally, we constructed an intracellular transport landscape that compares Marangoni-driven transport with thermal diffusion and motor-protein-mediated transport across cargo sizes. Together, our findings identify interfacial tension gradients as a previously unrecognized source of mechanical work in living cells and establish the Marangoni effect as a fundamental mechanism for the directed transport of mesoscale biomolecular condensates.

## Results

### Interfacial MEG-3 lowers the interfacial tension of PGL-3–RNA co-condensates

To determine whether MEG-3 directly regulates P granule interfacial properties, we reconstituted PGL-3–RNA co-condensates in vitro. Recombinant PGL-3–mCherry and MEG-3 were purified, and their RNA-binding activities were verified by electrophoretic mobility shift assays (EMSAs; Supplementary Fig. 2). Consistent with previous studies (*16, 17*), PGL-3–mCherry formed condensates with Alexa Fluor^TM^ 647-labelled *nos-2* CDS RNA (*nos-2*–AF647; **Methods**), establishing a minimal P granule model (Supplementary Fig. 3a). MEG-3–ATTO488 formed discrete clusters that selectively decorated the condensate interface (Fig. 1a).

**Fig. 1.**
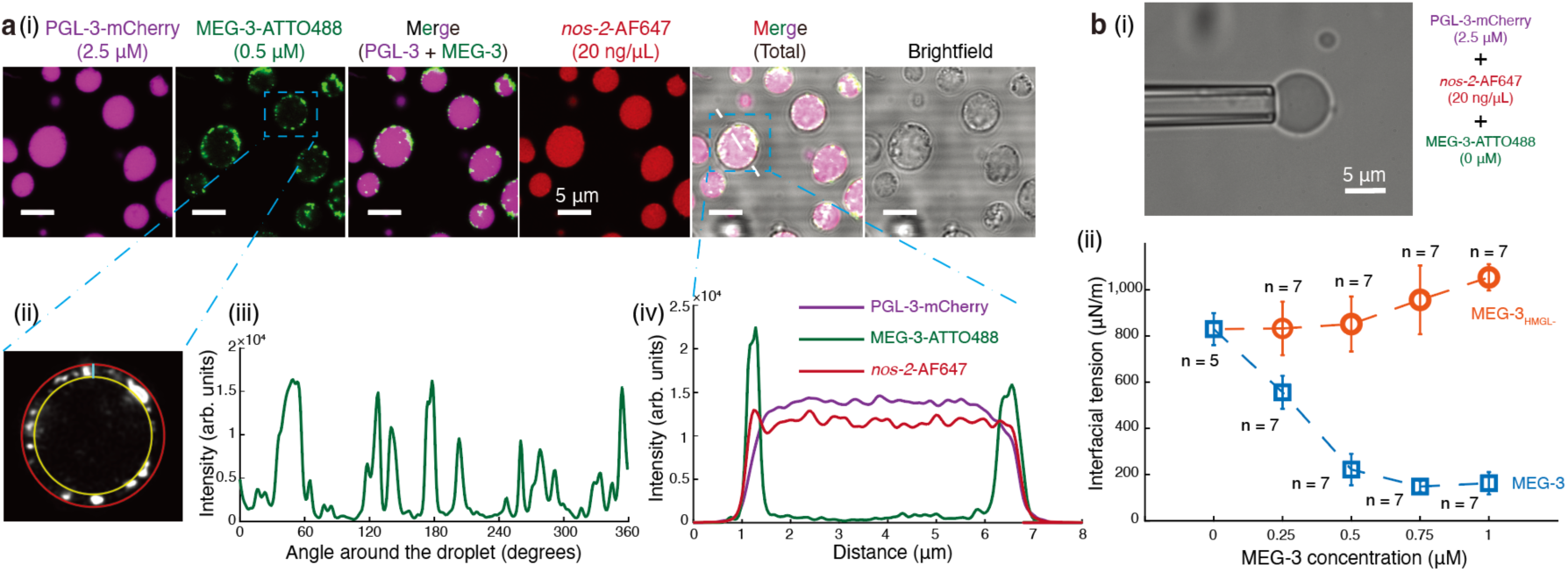
In vitro reconstitution of PGL-3–RNA co-condensates mimics P granules. (a) (i) 2.5 μM PGL-3–mCherry mixed with 20 ng/μL *nos-2*–AF647 RNA and 0.5 μM MEG-3-ATTO488; (ii)-(iii) Zoomed-in view of a single condensate, and angular distribution profile of MEG-3 intensity along the condensate periphery; (iv) Fluorescence intensity line profile across a representative condensate. Independent in vitro droplet assays were repeated three times (n = 3). (**b**) Micropipette aspiration assay. (i) Representative brightfield image of a reconstituted P granule aspirated into a micropipette; (ii) Interfacial tension (γ, μN/m) as a function of the concentration of MEG-3 (blue squares) and MEG-3_HMGL-_ (orange circles). The number n that independent experiments were repeated is labeled. Error bars, mean ± s.d. The working buffer used for these in vitro droplet assays was 25 mM HEPES (pH 7.5), 30 mM KCl, 90 mM NaCl, 1 mM DTT, 1.25% Glycerol.

Micropipette aspiration showed that increasing MEG-3 from 0 to 0.5 μM lowered interfacial tension approximately fourfold, from ∼800 to ∼200 μN/m (Fig. 1b and **Supplementary Information**). By contrast, the PGL-3-interaction-deficient mutant MEG-3_HMGL−_ (*17*) (Supplementary Fig. 2a(ii), b(ii), d, and 3b) did not measurably alter interfacial tension over the same concentration range but exhibited a modest upward trend at 0.5–1 μM concentration range instead, which may reflect altered interfacial interaction pattern between protein and RNA caused by weakened MEG-3–PGL-3 coupling (Fig. 1b and Supplementary Fig. 3b).

### MEG-3 gradients can drive directed transport of PGL-3–RNA co-condensates in vitro

To test whether spatially patterned MEG-3 drives condensate transport, we established a stable 0–0.5 μM MEG-3 gradient across a microfluidic channel containing PGL-3–RNA co-condensates (Fig. 2a and **Methods**). The gradient spanned several hundred micrometers, which was comparable to intrinsic MEG-3 gradient of *Caenorhabditis elegans* zygote (**Supplementary Information**) and remained stable for ∼1,000 seconds across various conditions, covering the relevant spatial and temporal scales (Supplementary Fig. 4). Condensates migrated persistently towards higher MEG-3 concentrations, whereas a MEG-3_HMGL−_ gradient produced no directional movement (Fig. 2b-c and Supplementary Movies 1-2).

**Fig. 2.**
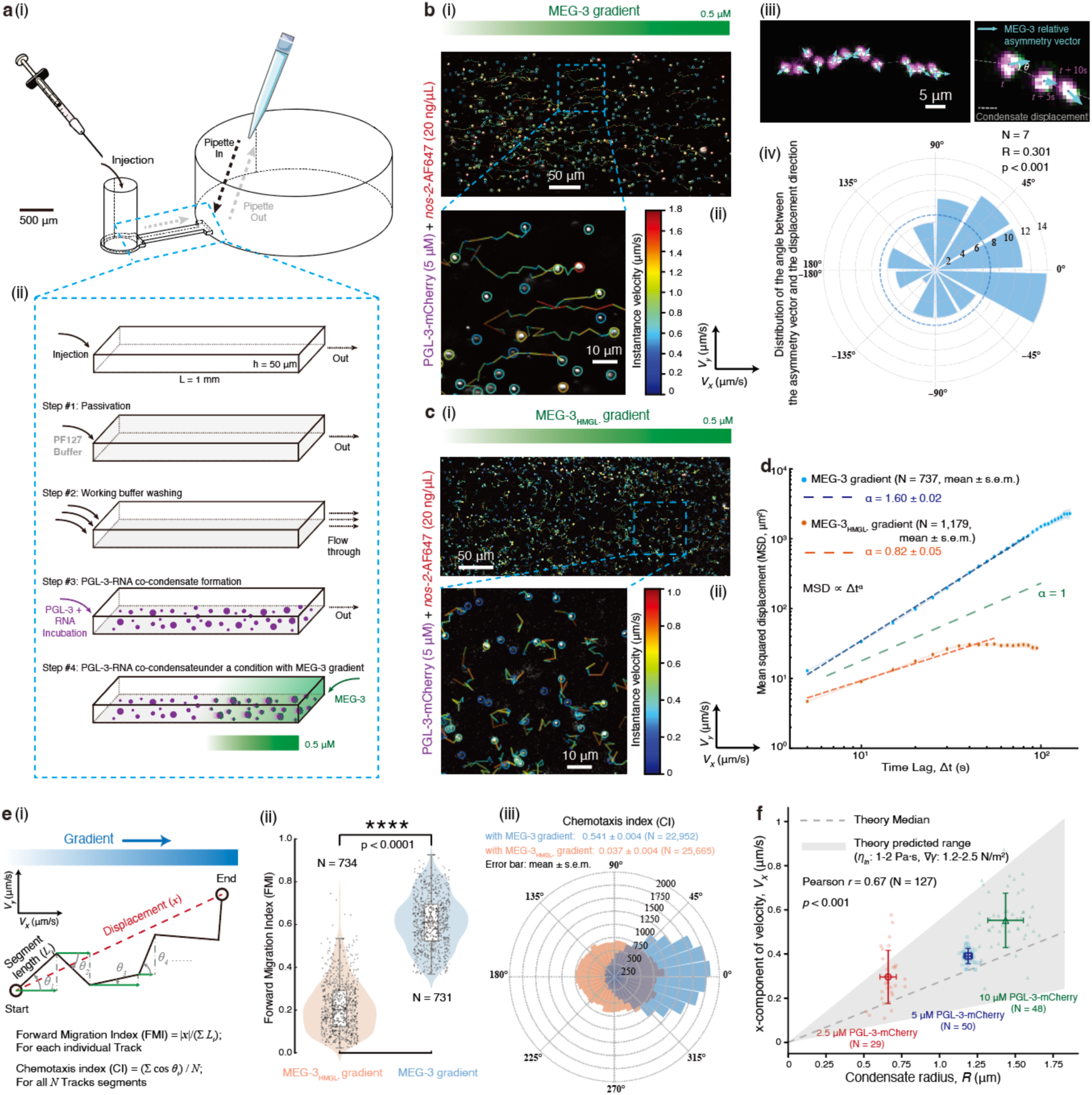
MEG-3 gradients induce directed transport of PGL-3–RNA co-condensates in vitro. (**a**) Schematic of our microfluidic device design (i); Experimental workflow (ii). (**b**) Directed motility of PGL-3–RNA co-condensates driven by the MEG-3 gradient. (i)-(ii) Condensate tracking trajectories of Supplementary Movie 1. (iii) Analyzing the relationship between the interfacial MEG-3 cluster distribution and the condensate displacement direction; (iv) Rose plot showing the distribution of the relative angle between the MEG-3 asymmetry vector and the displacement vector. 7 representative independent condensate tracking trajectories were analyzed. (**c**) Directed motility of PGL-3–RNA co-condensates driven by the MEG-3_HMGL-_ gradient. (i)-(ii) Condensate tracking trajectories of Supplementary Movie 2. (**d**) The MSD analyses of MEG-3 (737 condensates’ trajectories) and MEG-3_HMGL-_ gradient (1,179 condensates’ trajectories). Error bars, mean ± s.e.m. (**e**) (i) Schematic of FMI and CI. (ii) Boxplot of FMI distributions for condensates in MEG-3 (N = 731) and MEG-3_HMGL-_ (N = 734) gradients. For the boxplot, the black bar represents median. The bottom edge of the box represents 25^th^ percentiles, and the top is 75^th^ percentiles. Statistical significance was analyzed using two-sided Welch’s independent-samples t-test for two groups. p value: two-tailed; p value style: GP: 0.1234 (ns), 0.0332 (*), 0.0021 (**), 0.0002 (***), <0.0001 (****). Confidence level: 95%. (iii) Rose plot comparing the CI distributions. Error bars, mean ± s.e.m. (**f**) For theory, *V_x_* versus *R* were calculated via the YGB relation (Supplementary Table 1). The grey shaded area demarcates the parameter-free theoretical prediction range, and the dashed line represents the theoretical central value. For experimental data, *V_x_* shows a positive linear correlation with the radius *R* (Pearson r = 0.67, p < 0.001, N = 127). Colored markers represent experimental data obtained across varying PGL-3 concentrations. Error bars, mean ± s.d.

Mean-squared-displacement (MSD) analysis classified motion under the MEG-3 gradient as superdiffusive (α = 1.60 ± 0.02; N = 737, mean ± s.e.m.), while Motion under the MEG-3_HMGL−_ gradient was subdiffusive (α = 0.82 ± 0.05; N = 1,179, mean ± s.e.m.; Fig. 2d). Under the MEG-3 gradient, the mean velocity component parallel to the gradient (*V_x_*) was 0.37 ± 0.45 μm/s (N = 22,910, mean ± s.d.). The perpendicular component (*V_y_*) remained near zero (0.02 ± 0.41 μm/s; N = 22,910, mean ± s.d.). Both components were centered near zero under the MEG-3_HMGL−_ gradient (*V_x_* = 0.02 ± 0.32 and *V_y_* = 0.00 ± 0.32 μm/s, respectively; N = 57,948, mean ± s.d.) (Supplementary Fig. 5). The forward migration index (FMI) and chemotactic index (CI) independently confirmed directional bias only under the MEG-3 gradient (Fig. 2e). To test whether bulk MEG-3 gradient alone was sufficient, we pre-equilibrated condensates with 0.5 μM MEG-3 and imposed a 0.5–1 μM gradient. Condensates motion remained near-diffusive (α = 0.99 ± 0.02; N = 120, mean ± s.e.m.) and showed negligible directional bias (CI = 0.014 ± 0.010; N = 5,382, mean ± s.e.m.), consistent with the weak interfacial-tension response over this near-saturation range (Fig. 1b and Supplementary Fig. 6).

To relate directed transport to interfacial asymmetry, we compared the MEG-3 asymmetry vector with the frame-to-frame displacement vector for each condensate (Fig. 2b(iii)). The relative-angle distribution was non-uniform (mean resultant length R = 0.301; Rayleigh test, P < 0.001) and biased towards the MEG-3-enriched side (Fig. 2b(iv)). Thus, the imposed concentration gradient generated a local interfacial asymmetry on moving condensate, indicating that interfacial asymmetry tightly coupled to directed condensate motion.

### Condensate velocity positively scales with size

The directionality and interfacial asymmetry suggested that motion was propelled by Marangoni effect. Solving the Navier–Stokes equations of a spherical droplet under the assumptions of low Reynolds number and linear interfacial tension gradient yields the classical Young–Goldstein–Block (YGB) relation for the migration velocity of a Marangoni-driven droplet (*19*):

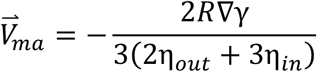

where *R* is the condensate radius, ∇γ is local interfacial tension gradient, η*_in_* is the condensate internal viscosity, and η*_out_* is the external viscosity of the surrounding medium.

At fixed ∇γ and viscosity, this relation predicts that migration velocity increases linearly with condensate radius. Combining the measured concentration dependence of interfacial tension, γ(*c*), with the imposed concentration gradient yielded ∇γ ∼1.2–2.5 N/m^2^ (Fig. 1b & Supplementary Table 1). Published measurements gave η*_in_* ∼1– 2 Pa·s (*16, 26*), whereas η*_out_* ∼10^-3^ Pa s was negligible by comparison. These values specify the parameter-free prediction of directed motion velocity (Fig. 2f).

In order to eliminate the impact of the Marangoni flow field crosstalk between adjacent condensates on individual motion, we varied the PGL-3 concentration to generate condensates with different size distributions, and calculated ensemble-averaged condensate characteristic size (*R*) and x-component velocity (*V_x_*) observed within each time point (**Supplementary Information**). Across various conditions, ensemble-averaged *V_x_*increased with characteristic radius (Pearson’s r = 0.67, P < 0.001, N = 127). The measured velocities fell within the predicted range (Fig. 2f). This quantitative agreement provides strong physical evidence that the long-range transport of reconstituted PGL-3–RNA co-condensates is governed by Marangoni effect.

### Alternative splicing isoform MEG-3 Δ16 reverses both interfacial response and transport direction

Inspection of *Caenorhabditis elegans* splice-junction annotations (*27*) identified an endogenous *meg-3* transcript lacking 48-nt within exon 2, generating a splice isoform that deletes 16 amino acids (residues 190–205; VSSVPVGFQKQENNSK; Supplementary Fig. 7a). This deleted segment lies within an intrinsically disordered region of MEG-3 (Supplementary Fig. 7b-c). We refer to this endogenous isoform as MEG-3 Δ16 and tested its properties with identical procedures (Supplementary Fig. 7d-e). Unlike MEG-3, recombinant MEG-3 Δ16 distributed uniformly along the condensate interface and partitioned substantially into the condensate interior (Fig. 3a and Supplementary Fig. 8). Increasing MEG-3 Δ16 from 0 to 0.5 μM raised interfacial tension from ∼800 to ∼1,050 μN m⁻¹ (Fig. 3b). MEG-3 Δ16 therefore reversed the sign of the interfacial tension response to concentration.

**Fig. 3.**
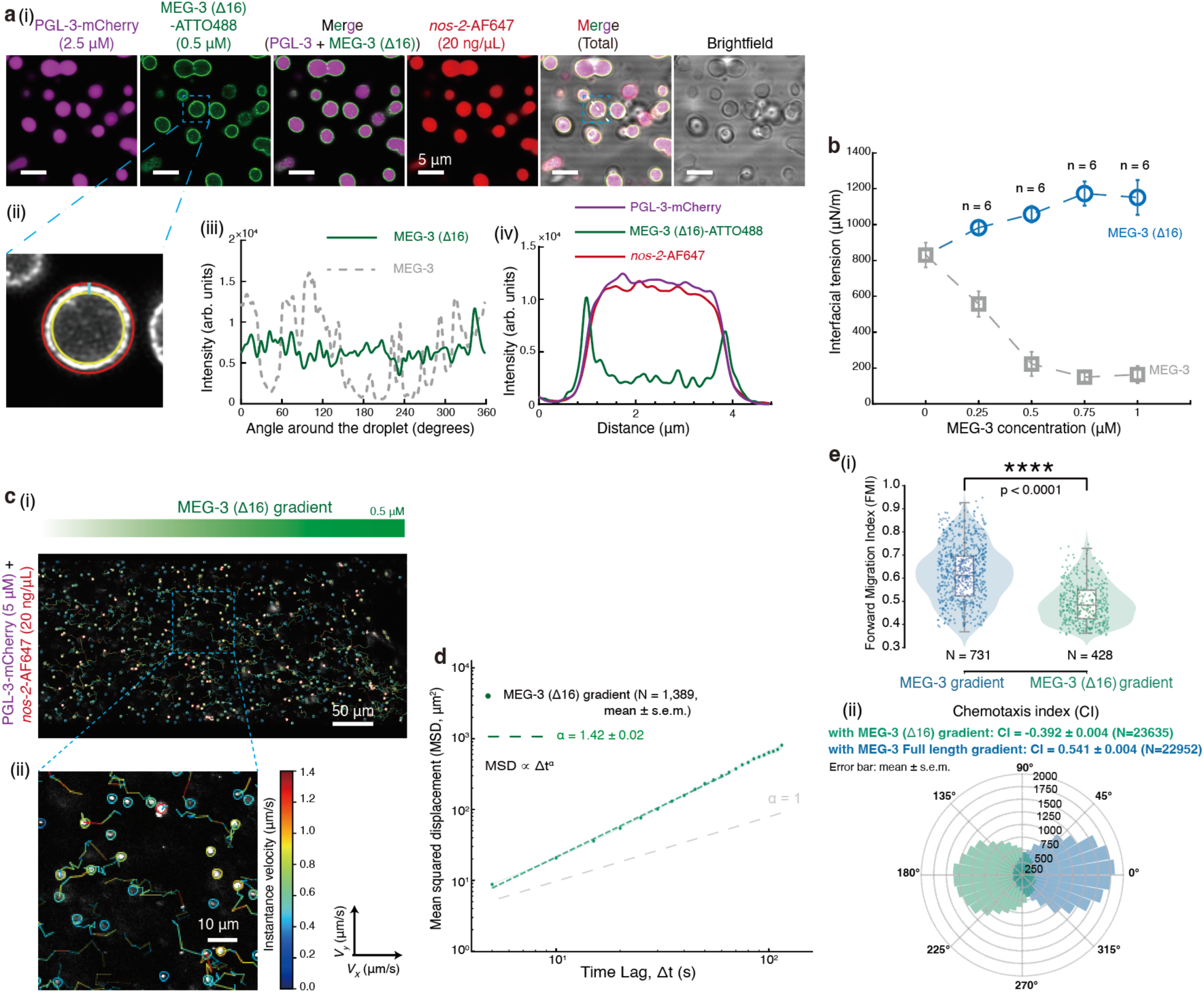
Alternative splicing isoform MEG-3 Δ16 reverses both interfacial response and transport direction. (**a**) (i) 2.5 μM PGL-3–mCherry mixed with 20 ng/μL *nos-2*–AF647 RNA and 0.5 μM MEG-3 Δ16-ATTO488; (ii)-(iii) Zoomed-in view of a single condensate, and angular distribution profile of MEG-3 Δ16 intensity along the condensate periphery; (iv) Fluorescence intensity line profile across a representative condensate. Independent in vitro droplet assays were repeated three times (n = 3). (**b**) Micropipette aspiration assay. Interfacial tension (γ, μN/m) as a function of the concentration of MEG-3 (gray squares) and MEG-3 Δ16 (blue circles). The number n that independent experiments were repeated is labeled. Error bars, mean ± s.d. (**c**) Directed motility of PGL-3–RNA co-condensates driven by the MEG-3 Δ16 gradient. (i)-(ii) Condensate tracking trajectories of Supplementary Movie 3. (**d**) The MSD analyses of MEG-3 Δ16 (1,389 condensates’ trajectories). Error bars, mean ± s.e.m. (**e**) (i) Boxplot of FMI distributions for condensates in MEG-3 (N = 731) and MEG-3 Δ16 (N = 428) gradients. For the boxplot, the black bar represents median. The bottom edge of the box represents 25^th^ percentiles, and the top is 75^th^ percentiles. Statistical significance was analyzed using two-sided Welch’s independent-samples t-test for two groups. p value: two-tailed; p value style: GP: 0.1234 (ns), 0.0332 (*), 0.0021 (**), 0.0002 (***), <0.0001 (****). Confidence level: 95%. (ii) Rose plot comparing the CI distributions. Error bars, mean ± s.e.m.

Consistent with this inverted response, a MEG-3 Δ16 gradient drove condensate migration opposite to that induced by MEG-3 (Fig. 3c, Supplementary Movie 3, and Supplementary Fig. 9). The mean axial and transverse velocities were *V_x_* = −0.23 ± 0.38 and *V_y_* = −0.02 ± 0.38 μm/s (N = 23,565, mean ± s.d.), respectively. Motion remained superdiffusive (α = 1.42 ± 0.02; N = 1,389, mean ± s.e.m.; Fig. 3d), and FMI and CI measurements confirmed directionality (Fig. 3e). Reversing the concentration– interfacial-tension relationship was therefore sufficient to reverse migration, directly linking transport direction to the interfacial tension gradient shaped by concentration gradient.

### P granule motion in embryos quantitatively agrees with Marangoni predictions

We next asked whether P granule motion in *Caenorhabditis elegans* embryos was quantitatively consistent with the Marangoni framework. Combining *γ*(*c*) with published MEG-3 concentrations and the anterior–posterior gradient yielded ∇γ ∼2.0–5.0 N/m^-2^ (Supplementary Table 1). The calculation used η*_in_* ∼2.5–10 Pa·s (*3, 16*), cytosolic basic viscosity *η_sol_* ∼0.01 Pa·s (*28, 29*), and cytoskeletal mesh size ξ ∼50 nm (*23, 30*) as previous studies described. We incorporated cytoplasmic resistance using the Brinkman–Darcy approximation for poro-viscoelastic drag (*31*), introducing an effective external viscosity, *η_out_*_,*eff*_ ∼ *η_sol_*(*R*/*ξ*)^2^, into the YGB relationship (Supplementary Table 1). This framework yielded a velocity range constrained by measurement or literature-derived physical quantities rather than fitting, enabling parameter-free prediction of P granule motility *in vivo* (Fig. 4a(iii)).

**Fig. 4.**
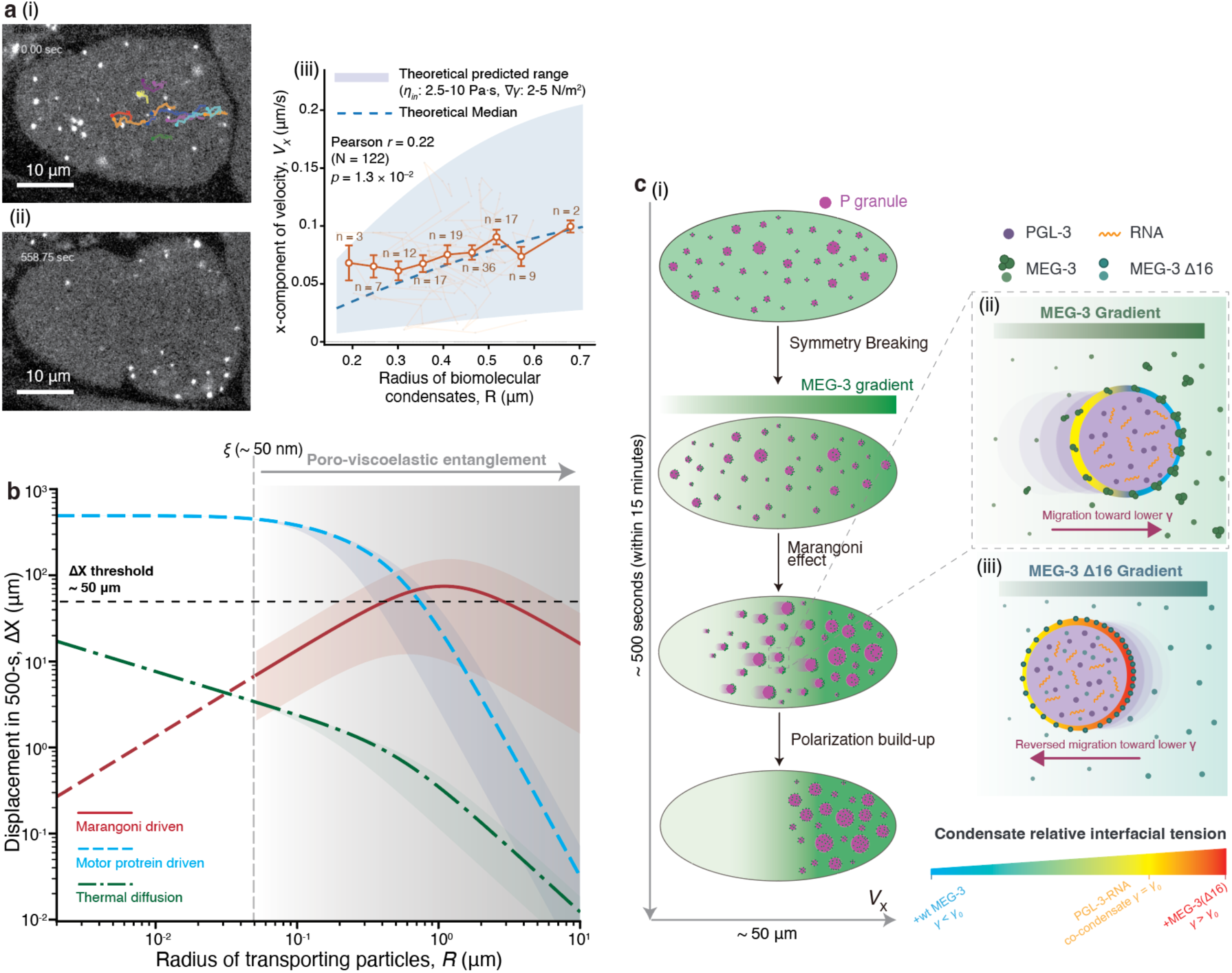
Marangoni effect as an existed highly efficient mechanism for biomolecular condensates intracellular transport. (**a**) In vivo tracking and kinematic analysis of P granules in a fertilized *Caenorhabditis elegans* one-cell zygote. (i)-(ii) Representative time-lapse confocal images of PGL-3::EGFP at the onset of symmetry breaking (t = 0.00 second, i) and after polarization (t = 558.75 seconds, ii). Overlaid tracks illustrate representative trajectories of directed motion toward the posterior pole. (iii) For theory, *V_x_* versus *R* were calculated via the YGB relation modified for the cytoplasmic poro-viscoelastic drag (Supplementary Table 1). The grey shaded area demarcates the parameter-free theoretical prediction range, and the blue dashed line represents the theoretical central value. For experimental data, the posterior-directed velocity *V_x_* exhibits a positive correlation with the radius *R* (Pearson r = 0.22, p = 1.3 × 10^-2^, N = 122). Data are binned by size. Error bars, mean ± s.e.m. (b) Theoretical transport landscape of micro-to-meso scale cargos within poro-viscoelastic cytoplasm environment. The predicted spatial displacement within a physiological 500 seconds time window (Δ*X*) is plotted against cargo radius (*R*) for three transport modes: thermal diffusion (green), molecular motor-driven transport (blue), and Marangoni-driven transport (red). The characteristic cytoskeletal mesh size ξ ∼50 nm. (**c**) (i) Schematic of Marangoni effect regulating the spatial organization of *Caenorhabditis elegans* P granules. (ii) for the condition with MEG-3 concentration gradient; (iii) for the condition with MEG-3 Δ16 concentration gradient.

Time-lapse imaging and particle tracking of PGL-3::EGFP (axIs1464) first zygote mitosis showed that approximately 60% of P granules underwent persistent anterior-to-posterior motion (Fig. 4a(i)-(ii), Supplementary Movie 4, and **Methods**). Approximately 60% of P granules underwent persistent anterior-to-posterior directional motion (Supplementary Fig. 1b). The tracked granules had a mean radius of 0.4 ± 0.1 μm (mean ± s.d.; N = 122). Posterior-directed velocity increased with granule radius (Pearson’s r = 0.22, P = 1.3 × 10^-2^, N = 122), and the measured velocities fell within the predicted range (Fig. 4a(iii)). Notably, previous works showed that series *meg-3* associated mutants, including *meg-3/4*, *mex-5/6*, MEG-3_HMGL-_/*meg-4*, MEG-3_Cterm_/*meg-4*, MEG-3_IDR_/*meg-4*; no matter applying perturbation on gradient establishment or P granule interface response disrupted P granule polarization (*3, 14, 16, 32, 33*). The magnitude and size dependence of P granule motion was therefore consistent with Marangoni propulsion in vivo.

### An intracellular transport landscape identifies a mesoscale regime for Marangoni-driven transport

Because the Marangoni effect naturally emerges from two fundamental features of intracellular organization—phase boundaries of biomolecular condensate and spatial biochemical heterogeneity, we next asked under what niche this mechanism becomes potentially effective for intracellular transport. To address this question, we constructed an intracellular transport landscape by comparing the characteristic transport distances achieved by diffusion, motor-driven transport, and Marangoni-driven transport as a function of cargo size over a fixed time interval (500 seconds; Fig. 4b and Supplementary Table 2). All three models incorporated poro-viscoelastic cytoplasmic resistance with a characteristic mesh size of ∼50 nm. Full force-balance derivations are provided in the **Supplementary Information**.

The resulting transport landscape reveals distinct physical regimes for intracellular cargo transport. Diffusion was effective for small molecules and nanoscale cargos but negligible for hundred-nanometer cargos, constrained by the randomness of direction (*25*). Motor-driven transport can directly move larger cargos but requires physical coupling to tracks and depends on adaptor availability, motor number and poro-viscoelastic cytoplasmic resistance. But its displacement decreased as *R*^−3^ for mesoscale cargo because resistance increased steeply with cargo size. Marangoni displacement increased with *R* when internal viscous dissipation dominated, while decreased as *R*^−1^ when external poro-viscous drag dominated (**Supplementary Information**). This produced a mesoscale regime favorable for Marangoni propulsion. For P granule-sized biomolecular condensates, the predicted displacement exceeded the ∼50-μm physiological distance. For cargos larger than ∼1 μm in radius, Marangoni-driven transport is predicted to surpass motor-driven transport in transport distance under the parameter set used here (Fig. 4b).

Together, these results define an intracellular transport landscape in which diffusion, molecular motors, and the Marangoni effect dominate complementary size regimes. Rather than replacing existing transport mechanisms, the Marangoni effect fills a hard-to-explain niche by providing an efficient mechanism for the long-range transport of mesoscale condensates that lack dedicated motor adaptors yet possess chemically tunable interfaces.

## Discussion

In this study, we identify the Marangoni effect as a mechanism by which an endogenous polarity gradient is converted into mechanical work at the interface and drive directed motion of a biomolecular condensate (Fig. 4c). In reconstituted PGL-3– RNA co-condensates, MEG-3 adsorbed to the condensate interface and reduced interfacial tension in a concentration-dependent manner. Consequently, a MEG-3 concentration gradient was sufficient to drive directional condensate migration. Weakening the MEG-3–PGL-3 interaction with the MEG-3_HMGL-_ mutant abolished both interfacial-tension regulation and directional transport, whereas the endogenous alternatively spliced isoform MEG-3 Δ16 reversed the concentration dependence of interfacial tension and correspondingly inverted the direction of condensate motion. In *Caenorhabditis elegans* embryos, both the magnitude and size dependence of P granule motion were quantitatively consistent with parameter-free predictions based on the endogenous MEG-3 gradient and independently measured material properties. Together, these findings suggest that P granule polarization includes a previously unrecognized Marangoni-driven transport component that operates alongside cytoplasmic flow and dissolution–recondensation. More broadly, our results establish a physical link between two ubiquitous features of living cells—phase boundaries of biomolecular condensates and spatial biochemical gradients, establishing a route of causality by which condensate emergent properties can contribute directly to cellular function. Over the past decade, numerous condensates have been identified in living cells and implicated in diverse cellular processes, yet their functional significance has often remained correlative. Our findings posit biomolecular condensates as functional active physical entities of intracellular organization and provides a framework understanding from the perspective of first principles.

An important question is how Marangoni-driven transport differs from typical phoretic transport. Phoretic mechanisms, including diffusiophoresis and osmophoresis, propel particles through slip or penetration flows generated within nanometre-scale interfacial boundary layers in response to chemical or osmotic gradients (*34–36*). By contrast, the Marangoni effect is driven by interfacial tension gradients acting directly along a fluid–fluid interface, producing tangential interfacial stresses that propel the entire condensate. Several observations distinguish the mechanism identified here from phoretic transport. First, Marangoni theory uniquely predicts that condensate velocity increases with condensate radius, a prediction that is quantitatively confirmed by our parameter-free analysis. Second, both the MEG-3_HMGL-_ mutant and the endogenous MEG-3 Δ16 isoform preserve the bulk MEG-3 concentration gradient while selectively altering the condensate interfacial response, thereby abolishing or reversing directional transport. Finally, the characteristic Marangoni velocities observed here (∼0.1–1 μm/s) exceed those typically reported for phoretic transport by orders of magnitude (**Supplementary Information**). Collectively, the key variable for transport is not the gradient alone, but the interfacial response function through which the spatial gradient is translated into ∇*γ* by the interface.

Our findings open several promising directions for future investigation. First, the Marangoni effect may represent a broadly conserved physical mechanism for intracellular spatial organization across diverse biological systems. Although diffusion tends to erase passive concentration differences, cells can sustain biochemical gradients as nonequilibrium reaction–diffusion states through spatially segregated modification cycles, differential binding and diffusion, localized turnover, or active transport (*37–40*). Biochemical gradients may therefore provide a widespread input for generating interfacial-tension gradients. This mechanism may therefore extend well beyond *Caenorhabditis elegans*. For example, *Drosophila oskar* RNP condensates regulated by their scaffold proteins (*41–43*), cleavage-furrow-localized zebrafish germ-plasm condensates in which Dnd1 controls RNA partitioning (*44, 45*), and Balbiani-body assembly composition turnover within *Xenopus* oocyte development (*46, 47*), all exhibit condensates spatial organization coupling with establishment of polarity cues and molecular gradients. These systems provide attractive experimental models in which gradient-dependent regulation of condensate interfacial properties could likewise generate Marangoni-driven transport and contribute to intracellular pattern formation.

Second, the Marangoni effect may represent a fundamentally cross-scale physical principle for biological self-organization. At larger length scales, analogous coupling between spatially patterned signals and interfacial tension gradients has been shown to generate global flows in multicellular systems, including cell aggregates and organoids(*48, 49*). These observations, together with our findings, raise the possibility that Marangoni-driven mechanics contribute to the organization of living matter across molecular, cellular, and tissue scales.

Third, the endogenous MEG-3 Δ16 splice isoform illustrates a novel paradigm for regulating condensate transport and function. Alternative splicing introduces only a subtle sequence variation within the intrinsically disordered region of MEG-3, yet reverses the sign of the concentration dependence of interfacial tension (*dγ*/*dc*; Fig. 3b), thereby inverting the direction of Marangoni-driven transport. More broadly, these findings raise the possibility that alternative splicing of intrinsically disordered proteins represents a potential regulatory dimension for remodeling the emergent properties like material properties, transport behavior, and biological functions of biomolecular condensates.

Another important open question is how condensate intrinsic rotation influences Marangoni-driven directionality. Because the driving polarity is imposed by an external gradient rather than fixed to the condensate body, intrinsic Brownian rotation need not abolish directed motion; instead, it should compete with the re-establishment of interfacial asymmetry. The resulting reorientation time scale is expected to reflect coupled timescales of Brownian translation and rotation, adsorption/desorption and partitioning of interfacial regulators, interfacial tension gradient relaxation and other kinetics. Their relative magnitudes determine macroscopic trajectory features, including transverse fluctuations, zigzagging, velocity autocorrelation and the lagged correlation with interfacial asymmetry. Future studies that tuning these kinetic parameters at molecular scale could establish predictive quantifiable links across scales and guide the design of condensate-targeting interfacial regulators with programmable cross-scale dynamics.

Taken together, our framework elevates condensate interfaces from uniform boundaries to programmable engines of intracellular organization. By tuning how local composition is translated into interfacial stress, cells may direct the motion, positioning and inheritance of membrane-free compartments across various concepts of biological process. Unveiling such response programs should reveal principles of active organization and enable rational control of transport in living systems.

## Author Contributions

H.Z. and Z.Q. conceived, designed, and supervised the project. Q.G. prepared biological samples, designed and conducted biochemical and microfluidics experiments, performed data analysis, built up the theoretical model, and wrote the paper. X.M. conducted *Caenorhabditis elegans* experiments. X.G. and J.L. conducted interfacial tension experiments. Z.Q. analyzed the data and wrote the paper. All authors commented on the manuscript.

## Acknowledgments

We thank microfluidic platform of the Center for Quantitative Biology, Peking University for PDMS microfluidic chip manufacturing support. We thank the Peking Nanofab for process support. We thank the contributions of the Engineering Research Center for Semiconductor Integrated Technology, Institute of Semiconductors, Chinese Academy of Sciences. We thank Dr. Yuansheng Cao (Tsinghua University) and Dr. Haoran Xu (Zhejiang University), and Dr. Cheng Li and other members of the Z.Q. laboratory for comments on the manuscript.

