## Supplemental methods for "Long-range transport of biomolecular condensates by Marangoni effect"

### Methods

#### Construction of bacterial expression plasmids

The expression plasmid pET28a-6×His::PGL-3–mCherry was described previously (*1*). The coding sequence for *Caenorhabditis elegans* MEG-3 (residues 1-862) was amplified from *Caenorhabditis elegans* cDNA. The MEG-3$_{HMGL-}$ mutant, containing the substitutions F705A, Y708A, Y713A, and N725A, was generated as described in a previous study (*2*). Both MEG-3, MEG-3$_{HMGL-}$ and MEG-3 Δ16 sequences were subcloned into the pET28a vector containing an N-terminal 6×His tag. All protein sequences used in this study are listed in **Supplementary Information**.

#### Protein purification

Purification of 6× His-tagged PGL-3::mCherry

The pET28a-6×His::PGL3::mCherry plasmid was transformed into E. coli strain Transetta (DE3), and cultured on LB plate at 37 °C for overnight. One single clone was picked and cultured in 20mL LB medium at 37 °C, 220 rpm for overnight. The culture of E. coli was diluted by the ratio of 1 : 100 to 2 L TB medium (+50 μg/mL Kanamycin) and cultured at 37 °C, 220 rpm. Until the OD600 was about 1.5, the bacteria was induced by 0.4 mM IPTG at 16 °C, 180 rpm for overnight to express the target protein. The E. coli cells were resuspended in binding buffer (50 mM Tris-Cl (pH 7.9), 500 mM NaCl, and 10 mM imidazole) sonicated with 1 mM PMSF and spun at 18,000 g for 30 minutes to pellet the debris. The supernatant lysates were purified on Ni-NTA beads. After extensive washing with binding buffer, the proteins were eluted with Elute Buffer (50 mM Tris-Cl (pH 7.9), 500 mM NaCl, and 500 mM imidazole), then further purified with a size column Superdex 200 increase GL 10/300 (GE Healthcare, USA) on an AKTA purifier (GE Healthcare) in PGL storage buffer (25 mM HEPES pH 7.5, 1 M

NaCl and 1 mM DTT). Purified protein solution was further concentrated with Amicon® Ultra Centrifugal Filter, 50 kDa MWCO (Merck Millipore) to a final concentration of ~10-20 mg/mL roughly estimated by Nanodrop spectrophotometer $A_{280}$ and calculated by Beer–Lambert law with the molar absorption coefficient of PGL-3::mCherry (79,760 $M^{-1}$ $cm^{-1}$).

<u>Purification of 6× His-tagged MEG-3 series proteins</u>

MEG-3 (aa 1-862), MEG-3$_{HMGL-}$ and MEG-3 Δ16 proteins were fused to an N-terminal 6×His tag in pET-28a then expressed and purified from inclusion bodies with denaturation protocols. The purification steps are same as PGL-3::mCherry purification before IPTG induce. The cultured bacteria induced at 1 mM IPTG at 16 °C, 180 rpm for overnight to express the target protein. Cells were resuspended in Lysis Buffer (25 mM HEPES (pH 8.0), 1,000 mM KCl, 10% (v/v) glycerol, 0.5% TritonX-100, and 0.5 mM TCEP), lysed by sonication under 1 mM PMSF, and spun at 25,000 g for 30 minutes. The supernatant was discarded and the pellet containing inclusion bodies was resuspended and washed in lysis buffer, briefly sonicated, and spun at 25,000 g, 30 minutes. The pellet was solubilized overnight at 4 °C in Buffer A with 6 M Urea. Spun the solubilized inclusion body at 25,000 g, 30 minutes and pass the supernatant over a Ni-NTA resin. Bound protein was washed with Wash Buffer (25 mM HEPES (pH 8.0), 1 M KCl, 25 mM Imidazole (pH 6), 10% (vol/vol) glycerol, 6 M urea, and 0.5 mM TCEP) and eluted in Elute Buffer (25 mM HEPES (pH 7.5), 1 M KCl, 250 mM Imidazole (pH 6), 10% (v/v) glycerol, 6 M urea, and 0.5 mM TCEP). Protein containing fractions were concentrated and further purified by size exclusion using a Superdex 200 increase GL 10/300 (GE Healthcare, USA) in Storage Buffer (25 mM HEPES (pH 7.5), 1 M KCl, 10% (v/v) glycerol, 6 M urea, and 1 mM DTT). Purified protein solution was further concentrated with Amicon® Ultra Centrifugal Filter, 50 kDa MWCO (Merck Millipore) to a final concentration of ~5–10 mg/mL roughly estimated by Nanodrop spectrophotometer $A_{280}$ and calculated Beer–Lambert law with the molar absorption coefficient of MEG-3/MEG-3 Δ16 (63,830 $M^{-1}\cdot cm^{-1}$) and MEG-$3_{HMGL-}$ (60,850 $M^{-1}\cdot cm^{-1}$).

All the target proteins were aliquoted and stored in −80 °C after quick freezing by liquid nitrogen. The concentrations of target proteins are further determined with BCA Protein Assay Kit P0012 (Beyontine, China) under dilution of each storage buffer without any reductant. All the experimental protein concentrations applied BCA results. Aliquots of peak elution fractions were run on ProteinEle® Precast Tris-Glycine Gel （4 – 20%）(Transgene, China), and stained with BeyoBlue™ Plus Coomassie Blue Super-Fast Staining Solution (Low Background) (Beyontine, China).

**Protein Labeling**

MEG-3 (aa 1-862), MEG-$3_{HMGL-}$ and MEG-3 Δ16 were labeled with reactive fluorophores ATTO 488 NHS-ester (Sigma-Aldrich). Fluorophores were mixed with protein by the molar ratio of 0.25–0.5 : 1 and incubate 15 minutes under room temperature. Reactions were quenched with 1/10 volume 1 M Tris-HCl (pH 7.4) to scavenge unreacted NHS-esters. Free fluorophore was eliminated by passage through three Amicon® Ultra Centrifugal Filter (30 kDa MWCO, 0.5 mL, Merck Millipore) into protein storage buffer. The concentration of fluorophore-labeled protein was determined using $A_{501}$ measured on a Nanodrop spectrophotometer and calculated with molar extinction coefficient of ATTO 488 (90,000 $M^{-1}\cdot cm^{-1}$).

**Electrophoretic mobility shift assay (EMSA)**

Binding reactions were performed by co-incubating purified proteins with 30 nucleotides (30-nt) Cy5-labeled RNA (5'-Cy5 – UUU CCC AAU CGU GGC GUG UCG GCC UCU UUU –3') at room temperature for 2 hours. The salt conditions in the reaction mixtures were maintained at 120 mM $K^+$ for the MEG-3 and MEG-$3_{HMGL-}$ assays, and 120 mM $Na^+$ for the PGL-3 assays. The working buffer contained 25 mM HEPES (pH 7.5), 120 mM KCl for MEG-3 and MEG-$3_{HMGL-}$, 120 mM NaCl for PGL-3–mCherry, 2 mM $MgCl_2$, 1 mM DTT, 5% Glycerol and 0.1 mg/ml BSA. Following incubation, the samples were resolved on a 6% native PAGE gel. Electrophoresis was conducted at 80 V for 90 minutes in a 4 °C cold room. The gel was subsequently visualized and recorded using a Typhoon gel imaging system.

**In vitro transcription of Alexa-Fluor 647 labeled *nos-2* CDS mRNA**

Alexa Fluor™ 647 labeled *nos-2* coding sequence (CDS) mRNA produced by *In Vitro* Transcription reaction (IVT) would be incubated together in reconstituted P granule as we referred previous studies (*2, 3*). The sequence of product RNA and template are presented in **Supplementary Information**. Template DNA for IVT was obtained by PCR amplification from Day 1 *Caenorhabditis elegans* cDNA. T7 promoter sequence (TAATACGACTCACTATAG) is added upstream *nos-2* CDS 5' end in the forward primer. The reaction is conducted in a 50 μL solution with ~1.5 μg *nos-2* CDS dsDNA template, 5 μL 10× reaction buffer (NEB), 200 units T7 RNA polymerase (NEB), 5 mM ATP, UTP, CTP, GTP for each (NEB), 50 nM Alexa Fluor™ 647-UTP (ThemoFisher), 50 units RNasin® Plus RNase Inhibitor (Promega), 10 mM DTT, 1 unit Yeast Pyrophosphatase, Inorganic (NEB). The mixture is incubated at 37 °C for 2 hours. Add 2 μL RNase free DNase I (Takara) to digest dsDNA template for 30 mins. Free NTPs and protein were removed by lithium chloride precipitation. RNAs were resuspended in water and stored at −80 °C. The integrity of RNA products was verified by 1% agarose gel electrophoresis and concentration was determined by $A_{260}$ with Nanodrop spectrophotometer and calculated by Beer–Lambert law.

**Preparation of PDMS Microfluidic Chips**

Microfabrication

Microfluidic channels were fabricated using standard soft lithography techniques. High-precision master molds were created by patterning SU-8 negative photoresist onto silicon wafers via photolithography. Polydimethylsiloxane (PDMS) base and curing agent (Sylgard 184, Dow Corning) were mixed at a 10 : 1 (w/w) ratio, degassed under vacuum to remove air bubbles, and poured over the SU-8 master. The replica was thermally cured at 65 °C for 4 hours. The cured PDMS slabs were cut and peeled from the master, and inlet/outlet ports were punched. The PDMS slabs were covalently bonded to cleaned glass coverslips following oxygen plasma treatment to seal the channels.

Passivation of Flow-Through Channels

For microfluidic channels with both inlets and outlets open, surface passivation was performed to prevent non-specific protein adsorption, following a protocol adapted from the Pluronic F127 coating procedure from (*4*) and described below. All solutions were introduced into the channels using clean glass syringes to minimize plasticizer contamination.

1. Etching: The channels were filled with 1 M KOH and incubated for 1 hour at room temperature to etch and activate the surface, followed by extensive rinsing with Milli-Q water to remove residual alkali.

2. Silanization: Sigmacote (Sigma-Aldrich) was injected into the channels to coat the surface. The reaction was allowed to proceed instantaneously, ensuring complete coverage. The channels were immediately flushed with isopropanol to remove excess silane and reaction byproducts, then dried thoroughly in the air.
3. Passivation: Prior to the experiment, the hydrophobized channels were filled with 0.5% (w/v) Pluronic F127 dissolved in the specific experimental buffer and incubated for >15 minutes. The channels were washed sequentially with the experimental buffer for at least 5 times.
4. Load experimental solution: Prepared experimental buffer was injected to channel and pre-incubated until flow within channel stayed still. Experimental buffer was pipetted into the inlet to generate solute gradient within channel for observation.

**Microscopy imaging**

<u>Droplet assay</u>

In vitro droplet imaging experiments were performed in Corning 384-well plates and images were acquired using a Leica TCS SP8 confocal microscope equipped with a 63× oil-immersion objective (NA 1.40). For the MEG-3 and PGL-3 co-condensation assays, the buffer conditions were maintained at 30 mM $K^+$ and 90 mM $Na^+$ and 20 ng/μL of AF647-labeled *nos-2* CDS RNA was included in the mixture. The working buffer used for these in vitro droplet assays was 25 mM HEPES (pH 7.5), 30 mM KCl, 90 mM NaCl, 1 mM DTT, 1.25% Glycerol. Standard excitation wavelengths corresponding to the respective fluorophores were applied. To precisely resolve the spatial distribution of MEG-3 signals on the surface of PGL-3 condensates, images were captured utilizing the Leica Lightning super-resolution mode.

<u>Microfluidic assay</u>

Following the device preparation as described in the previous section, microscopic imaging within the microfluidic channels was performed on the Leica TCS SP8 confocal microscope. Images were captured using a 20× air objective (NA 0.75). The ionic concentrations of the buffers utilized in the microfluidic assays were identical to those established for the droplet assays.

*In vivo* imaging of P granules dynamics

To monitor the dynamics of P granules in vivo, a *Caenorhabditis elegans* strain expressing PGL-3::EGFP (axIs1464, from (*5*)) was utilized. Continuous time-lapse imaging was performed using a Zeiss LSM880 confocal microscope. Specifically, the imaging focused on unlaid fertilized embryos undergoing cell division and polarization in utero. Time-lapse sequences were recorded in a single focal plane time lapse, using a time interval of 5–10 seconds per frame.

**Statistical test**

Statistical analyses were performed using Python with SciPy, NumPy, pandas, matplotlib and seaborn. Two-group comparisons of velocity distributions were performed using two-sided Mann–Whitney $U$ tests. Comparisons of Forward Migration Index (FMI) distributions were performed using two-sided Welch's independent-samples $t$-tests. Pearson's correlation coefficients were used to evaluate linear relationships between condensate size and migration velocity, and two-sided $P$-values were calculated. Directional distributions were evaluated using Rayleigh's test. Standardized effect sizes for pairwise velocity comparisons were calculated as Cohen's $d$ using the pooled standard deviation. Statistical significance was defined as $P < 0.05$, unless otherwise indicated.

### Statistical visualization and data presentation

Continuous data distributions were visualized using violin or raincloud plots combined with boxplots and jittered individual observations. In each boxplot, the central line denotes the median, the box limits represent the $25^{th}$ and $75^{th}$ percentiles, and whiskers indicate the plotted non-outlier range. Scatter plots show individual measurements or binned means, with error bars representing the standard deviation (s.d.) or standard error of the mean (s.e.m.), as indicated in the corresponding figure legends. Unless otherwise specified, n denotes independent experimental measurements, whereas N denotes analyzed trajectories or trajectory segments.
